# Repair of Noise-Induced Damage to Stereocilia F-actin Cores is Facilitated by XIRP2

**DOI:** 10.1101/2021.08.10.455815

**Authors:** E.L. Wagner, J.S. Im, M.I. Nakahata, T.E. Imbery, S. Li, D. Chen, Y. Noy, D.W. Archer, W. Xu, G. Hashisaki, K.B. Avraham, J.-B. Shin

## Abstract

Prolonged exposure to loud noise has been shown to affect inner ear sensory hair cells in a variety of deleterious manners, including damaging the stereocilia core. The damaged sites can be visualized as “gaps” in phalloidin staining of F-actin, and the enrichment of monomeric actin at these sites, along with an actin nucleator and crosslinker, suggests that localized remodeling occurs to repair the broken filaments. Herein we show that gaps in mouse auditory hair cells are largely repaired within one week of traumatic noise exposure through the incorporation of newly synthesized actin. Additionally, we report that XIRP2 is required for the repair process and facilitates the enrichment of monomeric *γ*-actin at gaps through its LIM domain-containing C-terminus. Our study describes a novel process by which hair cells can recover from sub-lethal hair bundle damage and which may contribute to recovery from temporary hearing threshold shifts and the prevention of age-related hearing loss.

## Introduction

Hair cells, the sensory receptors of the inner ear, are exposed to continuous mechanical stimulation from noise and head movement, which can, in some cases, be harmful to the sensitive structures of the cells, including the apical mechanosensitive apparatus known as the hair bundle (Wagner and Shin, 2019). The hair bundle, composed of filamentous (F)- actin-based stereocilia arranged in a staircase-like format, is deflected in response to mechanical stimulation, which causes the opening of mechanotransduction channels at stereocilia tips (Fettiplace, 2017; Hudspeth, 2005). Intense stimulation, such as from loud noise, can cause damage to the stereocilia F-actin cores, which is visible by transmission electron microscopy (TEM) as disorganization of the paracrystalline structure of negatively-stained actin filaments (Engström et al., 1983; Liberman, 1987; Liberman and Dodds, 1987; Tilney et al., 1982) or by confocal light microscopy as “gaps” in phalloidin-labeled F-actin (Avinash et al., 1993; Belyantseva et al., 2009). This damage is likely to decrease the rigidity of the bundle (Duncan and Saunders, 2000; Saunders et al., 1986), which could lead to reduced mechanotransduction (MET). In order to preserve hair cell function, an active repair mechanism would be needed to repair any damage to stereocilia F-actin. Passive repair through actin treadmilling is unlikely to be sufficient, because, unlike in most F-actin based structures, F-actin turnover in stereocilia is mostly restricted to a dynamic zone at stereocilia tips (Drummond et al., 2015; Narayanan et al., 2015; Zhang et al., 2012).

A previous study ((Avinash et al., 1993; Belyantseva et al., 2009) found that monomeric *β*- and *γ*- actin, as well as the actin-associated factors cofilin and espin, are enriched at phalloidin-negative gaps in stereocilia F-actin. The presence of monomeric actin, along with espin, which can crosslink actin filaments, and cofilin, which has actin severing activity and can nucleate F-actin assembly at high concentrations, led the authors to suggest that localized actin remodeling occurs at these sites to repair the damage. It was also shown that inner hair cell stereocilia in mice lacking *γ*-actin develop gaps in the absence of overstimulation. Therefore *γ*-actin is likely important for the repair of gaps or in the prevention of their formation. However, they did not conclusively show that this damage can be repaired or how repair might occur.

Herein we propose a role for XIRP2 in the repair process. XIRP2 expression is enriched in hair cells where it colocalizes with F-actin-based structures (Francis et al., 2015; Scheffer et al., 2015). XIRP2 is expressed in two main isoforms: a long isoform that includes a known F-actin binding domain, and a short isoform that is lacking the known F-actin binding domain but includes a C-terminal tail with a predicted LIM domain. The long isoform primarily localizes to the cuticular plate and cell junctions, while the short isoform is enriched in stereocilia (Francis et al., 2015). XIRP2 knockout mice have normal hair bundle development, but F-actin core disruption is visible as early as P7 (Scheffer et al., 2015) and stereocilia degeneration is detectable by P12, leading to hearing loss by seven weeks of age (Francis et al., 2015).

Many LIM domain-containing proteins are associated with cytoskeletal processes, including the repair of stress fiber strain sites (SFSS) (Smith et al., 2013, 2010). Similar to the gaps in stereocilia actin cores, SFSS appear as dimmed sections in fluorescently labelled F-actin stress fibers. Zyxin and paxillin are recruited to these SFSS in a LIM-domain dependent manner and subsequently recruit an actin nucleator (VASP) and an actin crosslinker (*α*-actinin) to the SFSS to repair the damage and prevent a catastrophic break of the stress fiber. Additionally, some LIM domains have been shown to be mechanoresponsive, binding directly to tensed F-actin (Sun et al., 2020), which may be present at SFSS and gaps in stereocilia cores.

In this study, we demonstrate that the number of noise-induced gaps in murine hair cell stereocilia actin cores decreases to baseline levels within one week of exposure and that newly synthesized actin is incorporated into the stereocilia core in this time frame, suggesting that the damage is repaired through localized F-actin remodeling. We also show that XIRP2 is recruited to gaps through its LIM domain-containing C-terminus, and is required for their repair, likely through the recruitment of monomeric actin. Gap repair may also contribute to the prevention of age-related hearing loss and recovery from temporary hearing threshold shifts, as *Xirp2* knockout mice develop progressive hearing loss and have worse permanent hearing loss than wild types following traumatic noise exposure.

## Results

### Exposure to loud noise causes reversible damage to the stereocilia F-actin core

As was previously reported in guinea pigs (Avinash et al., 1993; Belyantseva et al., 2009), exposure to prolonged loud noise causes damage to the core of sensory hair cell stereocilia (**Fig. 1A-B**), visualized as “gaps” in phalloidin labeling of F-actin (**Fig.1C**), which are most easily observed in the tallest row stereocilia of inner hair cells (IHCs). In adult mice, gaps are present at a low rate (0.08±0.023 gaps per IHC) in mice unexposed to prolonged loud noise, but following 1 hour of 120 dB broadband (1-16kHz) noise, this increases to 0.35±0.085 gaps per IHC (**, p=0.005) (**Fig. 1D**). We chose to focus on IHCs in this study both because noise-induced damage appears to be more severe in IHC bundles than in OHCs (Liberman, 1987) and because of their relative ease of visualization with light microscopy compared to OHCs.

**Figure 1.**
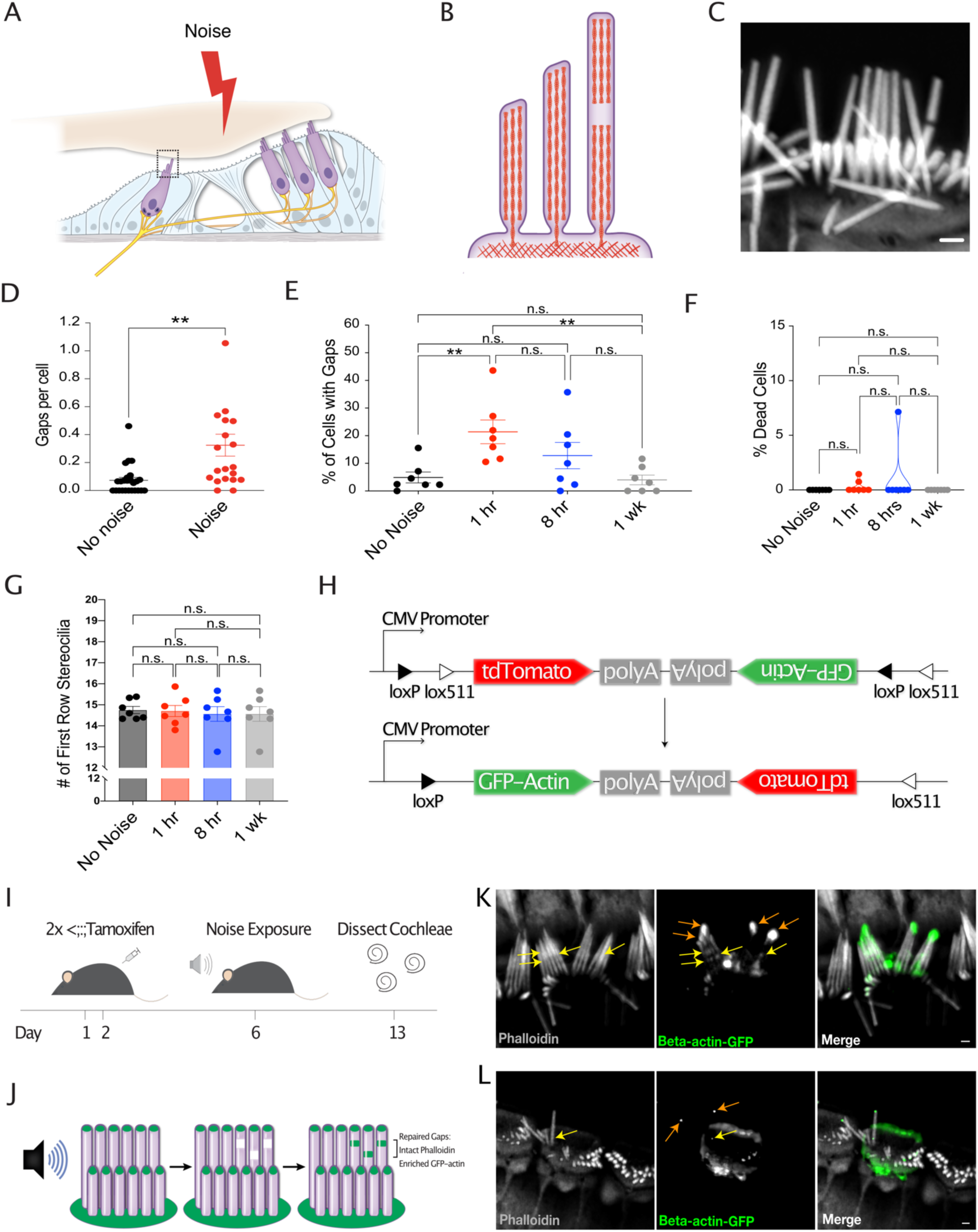
Intense noise exposure causes reversible damage to the F-actin cores of stereocilia. **(A)** Cartoon showing cross section of the organ of Corti. **(B)** Cartoon depicting side view of an inner hair cell (IHC). **(C)** Cartoon representing gap in IHC stereocilia F-actin following noise exposure and representative image. Gap in phalloidin staining indicated by arrow. Scale bar: 1μm. **(D)** Increased number of phalloidin-negative gaps in IHC stereocilia F-actin per cell following noise exposure (**, p=0.0053). No Noise: n=25 organs of Corti, 13 mice; Noise: n=20 organs of Corti, 10 mice. **(E)** Percentage of cells with gaps initially increases 1 hour following noise (p=0.004) but then decreases (p=0.003) to levels not significantly different than in unexposed mice (p=0.74). n=7 organs of Corti, 4 mice per group. **(F)** Percentage of dead inner hair cells per cochlea does not significantly change within 1 week of noise exposure (No Noise vs 1 hr, n.s., p=0.174, No Noise vs 8hrs, n.s., p=0.337, No Noise vs 1 week, all values 0). n= 7 organs of Corti, 4 mice per group. **(G)** Number of tallest row stereocilia per hair cell does not significantly change at any measured point within 1 week of noise exposure. No Noise vs 1 hr-n.s., p= 0.888, No Noise vs 8hrs, n.s., p=0.641, No Noise vs 1 week, n.s., p=0.854. **(H)** Diagram of Cre-mediated inversion in *FLEx-β-actin-GFP* mice following tamoxifen injection. Expression of tdTomato is turned off and GFP-*β*-actin expression is turned on. **(I)** Experimental schematic for the observation of the localization of newly synthesized GFP-tagged *β*-actin. Mice are injected on days 1 and 2 with tamoxifen and exposed to noise on day 6. Cochleae were dissected and processed on day 13. **(J)** Cartoon demonstrating the expected localization of GFP-tagged *β*-actin in repaired gaps. **(K-L)** Representative examples from >8 experiments of likely repaired gaps. Yellow arrows point to sites of enriched GFP-tagged *β*-actin along stereocilia length with intact phalloidin staining in a Cre-recombined cell following one week of recovery from noise exposure. Orange arrows indicate GFP-labeled stereocilia tips. Due to low recombination rates, the surrounding cells do not express GFP-*β*-actin.. All error bars represent the standard error of the mean (SEM).

The enrichment of monomeric actin and an actin nucleator and crosslinker at gaps suggested that new F-actin polymerization might be occurring to repair the damage (Belyantseva et al., 2009), but direct evidence of this was lacking. In order to determine whether gaps are repaired, we quantified the percent of inner hair cells with gaps in a time course following traumatic noise exposure (4hrs, 120 dB broadband noise). Following an initial increase in the percent of IHCs with gaps from 4.89±1.97% to 21.39±4.29% in mice 1 hour following noise, the percent of IHCs with gaps decreased to 12.80±4.76% by 8 hours post noise (n.s., p=0.20) and to 4.00±1.76% by 1 week post noise (**, p=0.003). The percent of IHCs with gaps 1 week following noise exposure was no longer significantly different from the baseline level (n.s., p=0.74), suggesting that noise-induced gaps are largely repaired within this timeframe (**Fig. 1E**).

To address the possibility that the decreased percentage of IHCs with gaps was not due to the death of damaged cells, we quantified the amount of IHC death at each timepoint before and after noise exposure. There was no significant increase in the number of dead IHCs at any time point after noise (No Noise vs 1 hr, p=0.174, No Noise vs 8hrs, p=0.337, No Noise vs 1 week, all values 0) (**Fig. 1F**), so it is unlikely that the death of damaged hair cells contributed to the observed decrease in cells with gaps. Another potential explanation for the decrease in the percent of IHCs with gaps was the shedding of damaged stereocilia. However, we also did not observe any significant change in IHC tallest row stereocilia number following noise exposure (No Noise vs 1 hr, p=0.888, No Noise vs 8hrs, p=0.641, No Noise vs 1 week, p=0.854) (**Fig. 1G**), consistent with repair being the primary cause of the decrease in the percent of IHCs with gaps.

We hypothesized that gaps repaired by localized remodeling would incorporate newly synthesized actin. Therefore, as an alternative approach to demonstrate the repair of gaps, we looked at the localization of actin synthesized following noise exposure, using *FLEx β-actin-GFP* mice crossed to a tamoxifen-inducible Cre line (*Ubc-Cre^ERT2^*), in which, following Cre recombination, *β-actin-GFP* expression is turned on (**Fig. 1H**). Four days following tamoxifen injection, to provide time for maximal Cre recombination, we exposed *FLEx-β-actin-GFP+;Ubc-Cre^ERT2^+* mice to traumatic noise (4 hrs, 120 dB broadband) and allowed the mice to recover for 1 week (**Fig. 1I**), during which time gaps were repaired in our previous experiment. Due to minimal actin turnover in stereocilia, when unexposed to noise, *β*-actin-GFP is localized only to stereocilia tips in these mice (Narayanan et al., 2015). However, if newly synthesized *β*-actin is incorporated into gaps during repair, we would expect to see patches of *β*-actin-GFP at discrete sites along the length of stereocilia with intact phalloidin staining in addition to the staining at the stereocilia tips (**Fig. 1J**). Consistent with this, we see areas of enrichment of newly synthesized *β*-actin in stereocilia following noise exposure which are likely sites of repair (**Fig. 1K-L, yellow arrows**). Due to low Cre recombination rates (∼10% of IHCs) and dim *β*-actin-GFP fluorescence, quantification of the occurrence of these “repaired gaps” was not possible, but their presence supports the complementary evidence that noise-induced stereocilia core damage is repaired.

### XIRP2 is enriched at gaps and is required for their repair

Monomeric *β*- and *γ*-actin and several actin-associated factors have been shown to be enriched at gaps, where they likely contribute to the repair process (Belyantseva et al., 2009). In addition to these, we observe the enrichment of XIRP2 immunostaining at gaps (**Fig. 2A-B**). This enrichment is observed both in naturally occurring gaps in murine auditory inner hair cells of the cochlea (**Fig. 2A**) and vestibular hair cells of the utricle (**Fig. 2B**), including gaps which appear to span across several neighboring stereocilia (**Fig. 2B**). These adjacent gaps may occur as a consequence of the interconnectedness of the hair bundle, with a mechanical break in one stereocilium destabilizing its neighbors in a cascading manner.

**Figure 2.**
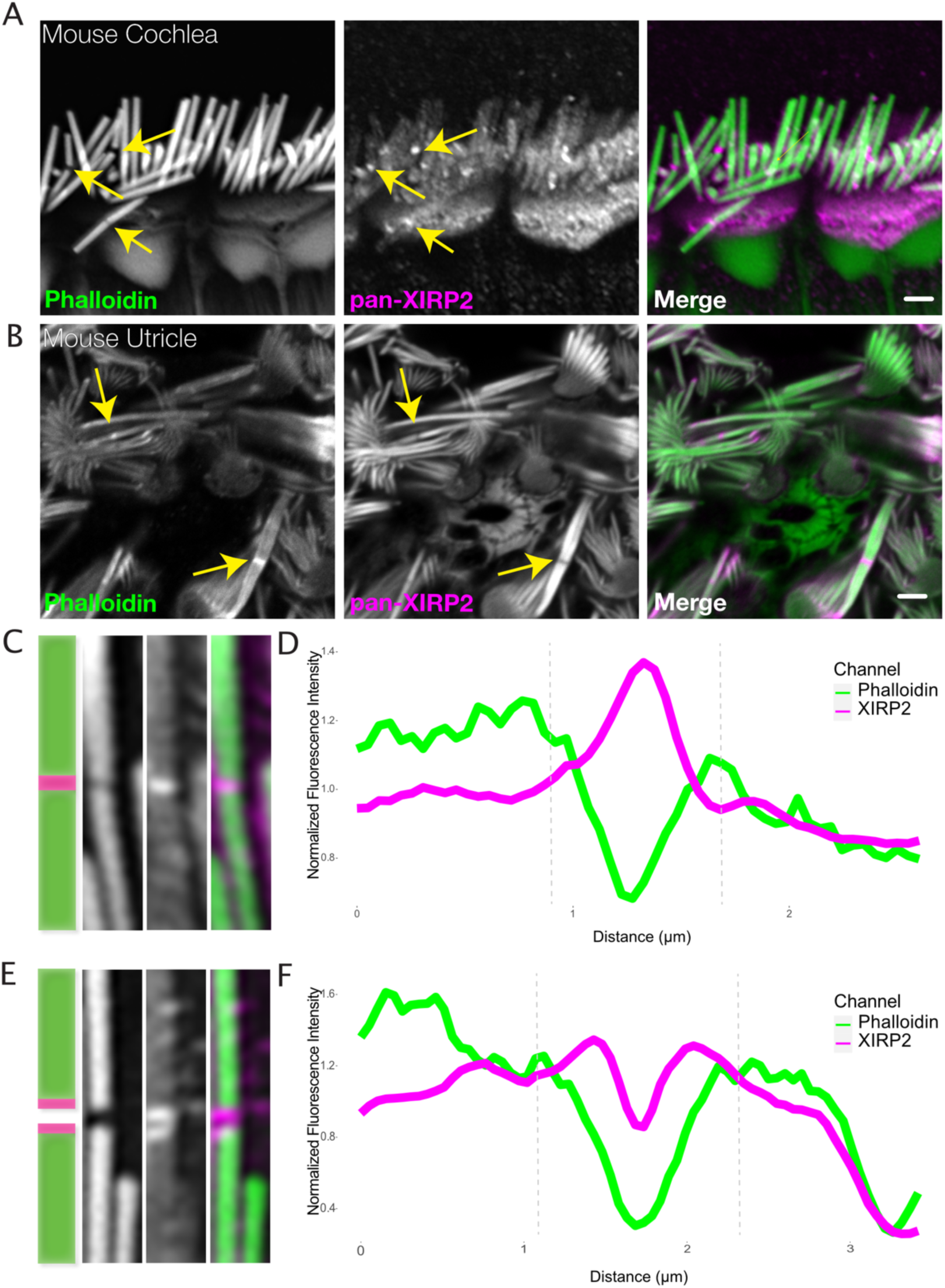
XIRP2 is enriched at gaps in stereocilia F-actin cores. **(A)** XIRP2 immunostaining is enriched at noise-induced gaps in phalloidin staining in IHCs (yellow arrows). **(B)** XIRP2 immunostaining is enriched at naturally-occurring phalloidin-negative gaps in utricle hair cells (yellow arrows). **(C)** Cartoon and representative image (1×5μm) of XIRP2 enrichment throughout length of gap in phalloidin signal. **(D)** Line scan of fluorescence intensity in phalloidin (green) and XIRP2 (magenta) channels along the length of a gap in which XIRP2 is enriched throughout the gap. **(E)** Cartoon and representative image (1×5μm) of XIRP2 enrichment at the edges of a gap. **(F)** Line scan of fluorescence intensity in phalloidin (green) and XIRP2 (magenta) channels in which XIRP2 is only enriched at the edges of the gap. Images are representative of >10 experiments.

A representative line scan plotting the fluorescence of XIRP2 shows an approximately 1.4-fold increase in XIRP2 intensity in the center of a phalloidin-negative gap compared to the surrounding area of the stereocilia core (**Fig. 2C-D**). Occasionally, in larger phalloidin-negative gaps, we observe a gap in the XIRP2 staining as well, with enrichment only at the gap edges (**Fig. 2E-F**). XIRP2 enrichment is present both in gaps induced by loud noise exposure (**Fig. 3B**) and in gaps occurring in genetic models of hair bundle degeneration (Goodyear et al., 2012; Krey et al., 2018), including *Ptprq* (**Fig. 3C**) and *Elmod1* knockout mice (**Fig. 3D**). Additionally, we confirmed that this staining pattern is present in human hair cells, as well (**Fig. 3E**). To ensure that the enrichment of XIRP2 immunostaining at these sites was not an artifact of antibody staining, we found that the XIRP2 was no longer enriched at gaps in *Xirp2* knockout mice (Francis et al., 2015) (**Fig. 3F**). Moreover, we see XIRP2 enrichment at gaps using a second XIRP2 antibody specific for the short isoform, which is localized to the hair bundle (**Fig. 3G**), but not an antibody specific for the long isoform, which is primarily localized to other F-actin-based hair cell structures (Francis et al., 2015) (**Fig. 3H**). A gene map in **Fig. 3A** indicates the position of the epitopes for the pan-XIRP2 and isoform specific antibodies.

**Figure 3.**
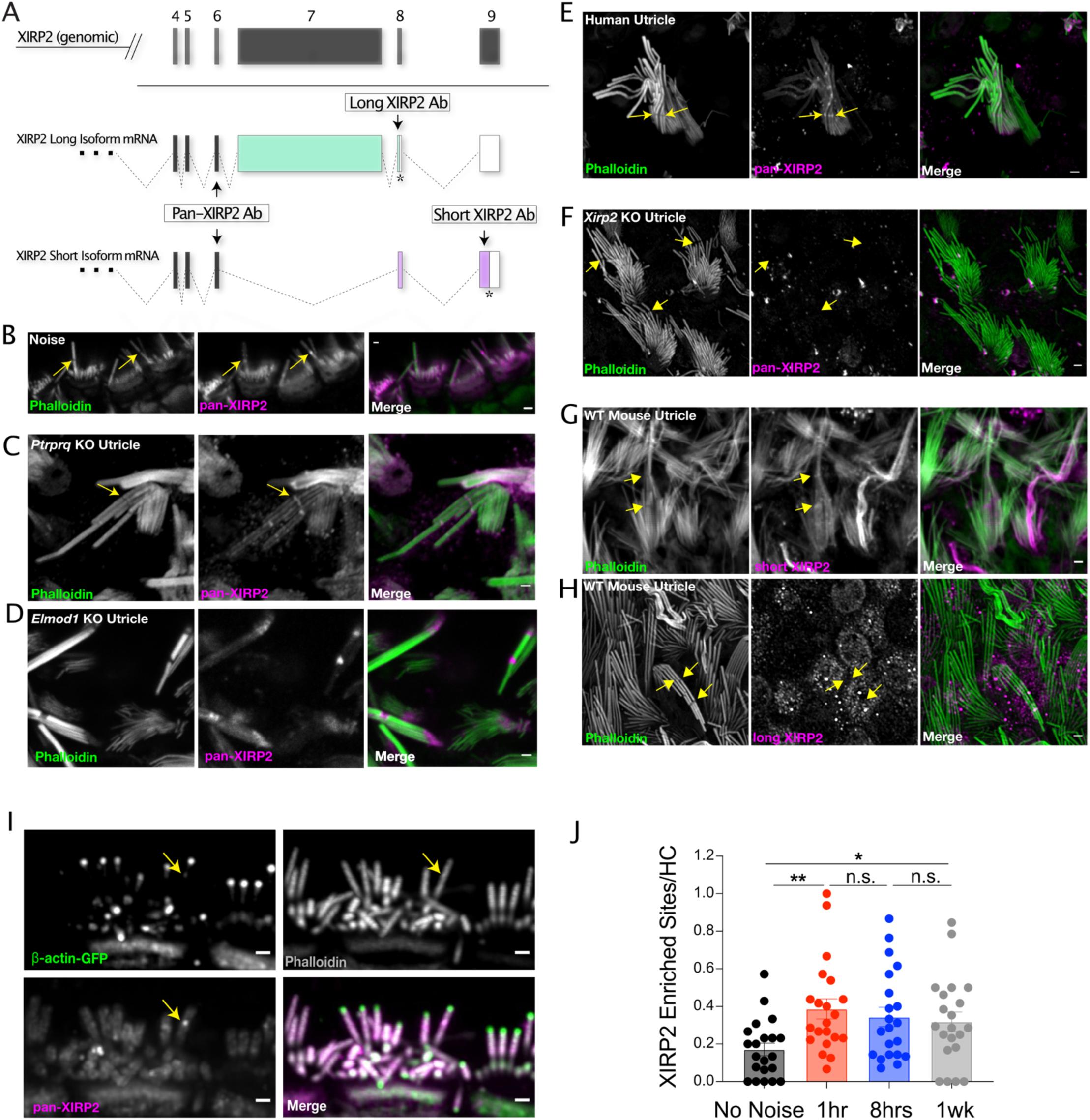
The short isoform of XIRP2 is enriched at gaps and remains there after repair. **(A)** Diagram of *Xirp2* gene structure and isoforms indicating the positions encoding the epitopes recognized by antibodies used in the figure. **(B)** XIRP2 enrichment at gaps (yellow arrows) in IHCs induced by overexposure to noise (120dB broadband (1-22kHz) for 4hrs). **(C)** XIRP2 enrichment at gaps in a utricle hair cell in a P25 *Ptrpq* knockout mice. **(D)** XIRP2 enrichment in gaps (yellow arrows) in IHCs in a *P20 Elmod1* knockout mouse. **(E)** XIRP2 enrichment at gaps (yellow arrows) in a human utricle hair cell. **(F)** XIRP2 staining is absent in *Xirp2* knockout utricle hair cells. **(G)** Short XIRP2 is enriched at gaps (yellow arrows). **(H)** Long XIRP2 is excluded from the hair bundle and not present in gaps (yellow arrows). **(I)** XIRP2 enrichment is colocalized with sites of enriched synthesized *β*-actin synthesized after noise exposure that likely represent repaired gaps (yellow arrow). **(J)** The number of XIRP2 enriched sites in stereocilia increases following noise exposure ((No Noise vs 1 hr, **, p=0.002, 1hr vs 8hrs, n.s., p=0.569, 8hrs vs 1 week, n.s., p=0.731, No Noise vs 1 week, *, p=0.021). n= 21 images; 7 organs of Corti; 4 mice per group. All scale bars are 1μm. Error bars represent SEM. Images are representative of >3 experiments.

Interestingly, XIRP2 appears to remain enriched at repaired gaps, as we see colocalization of XIRP2 enrichment at sites labeled by enriched *β*-actin synthesized after noise exposure (**Fig. 3I**). Consistent with this, we also see an enrichment of XIRP2-enriched sites in stereocilia cores, which remains significantly increased 1 week following noise exposure (No Noise vs 1 hr, **, p=0.002; 1hr vs 8hrs, n.s., p=0.569; 8hrs vs 1 week, n.s., p=0.732; No Noise vs 1 week, *, p=0.021) (**Fig. 3J**), unlike the number of gaps, which decreases during this time period.

The presence of XIRP2 in gaps suggested that it was playing a role in their repair. To address this possibility, we evaluated the capacity for repair in *Xirp2* knockout mice. We first examined the gaps in vestibular and auditory hair cells that occur in the absence of exposure to loud noise. In *Xirp2* knockout mice, abundant gaps were observed in both vestibular hair cells in the utricle (**Fig. 4A**) and in IHCs (**Fig. 4C**). At postnatal day 6, *Xirp2* knockout mice had 4.33-fold more gaps per utricle hair cell than in wild type mice (***, p<0.0001) (**Fig. 4B**). By postnatal day 20, there were also significantly more gaps per IHC in *Xirp2* knockout mice compared to WT (**, p=0.0015) (**Fig. 4D**). These results, concordant with a previous report of F-actin filament disorganization in a different *Xirp2*-null mouse line (Scheffer et al., 2015), suggest that gaps may build up over time in the absence of XIRP2.

**Figure 4.**
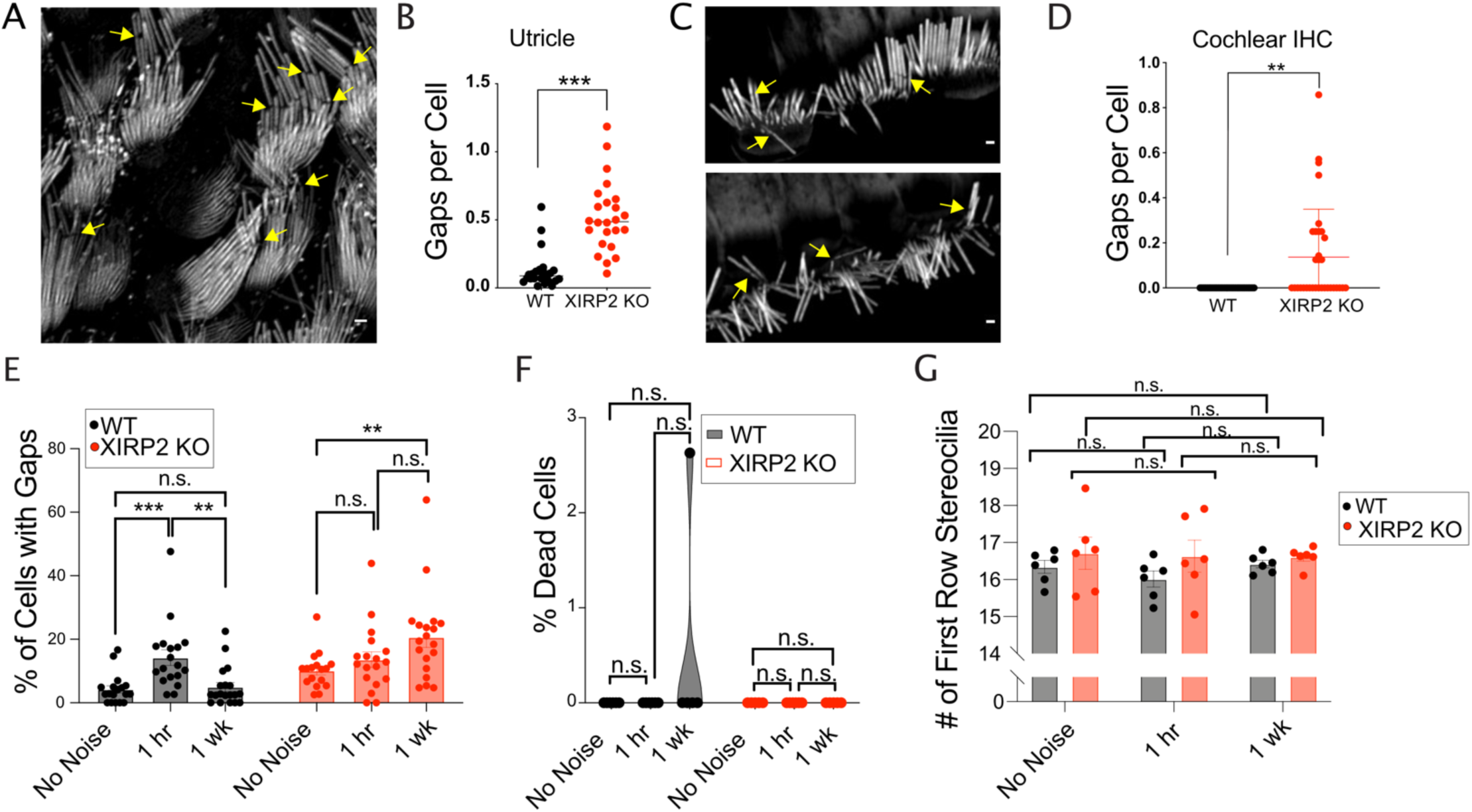
XIRP2 is required for the repair of gaps. **(A)** Gaps (yellow arrows) in stereocilia F-actin in utricle hair cells from *P6 Xirp2* knockout mice. **(B)** There are significantly more gaps per hair cell in P6 *Xirp2* knockout mice than in age-matched WT mice (***, p<0.0001). n=24 images, 6 utricles, 3 mice per group. **(C)** Gaps (yellow arrows in stereocilia F-actin in cochlear IHCs from P20 *Xirp2* knockout mice. **(D)** There are significantly more gaps per hair cell in P20 *Xirp2* knockout mice than in age-matched WT mice (**, p=0.002). n=24 images, 6 utricles, 3 mice per group. **(E)** Percentage of cells with gaps decreases to before noise levels within 1 week after initial increase in WT mice, but in *Xirp2* knockout mice, the percentage of cells with gaps actually increases during this time period. WT: No noise vs 1hr - ***, p<0.001, 1 hr vs 1 wk - **, p=0.002, No noise vs 1 wk - n.s., p=0.68; *Xirp2* knockout (KO): No noise vs 1hr- n.s., p=0.45, 1 hr vs 1wk – n.s., p=0.09, No noise vs 1wk - **, p=0.003. n= 18 organs of Corti, 9 mice per group. **(F)** Percentage of dead inner hair cells per cochlea does not significantly change within 1 week of noise exposure in WT or *Xirp2* knockout mice. WT: No Noise vs 1 hr- all values 0, No Noise vs 1 week, n.s., p=0.29; *Xirp2* KO: No Noise vs 1 hr – all values 0, No Noise vs 1 week – all values 0. n= 6 organs of Corti, 3 mice per group. **(G)** Number of tallest row stereocilia per hair cell does not significantly change within 1 week of noise exposure in WT or *Xirp2* knockout mice. WT: No Noise vs 1 hr- n.s., p= 0.263, No Noise vs 1 week, n.s., p=0.715; *Xirp2 KO*: No Noise vs 1 hr – n.s., p=0.901, No Noise vs 1 week – n.s., p=0.820. n= 6 organs of Corti, 3 mice per group. All scale bars are 1μm. Error bars represent SEM. Images are representative of >3 experiments.

To more directly assess the necessity of XIRP2 in the gap repair, we quantified the percent of IHCs with gaps in a time course following noise exposure (4hrs, 120 dB broadband) in wild type and *Xirp2* knockout mice. In wild type mice, after an initial increase 1 hour post noise (***, p<0.001), the percent of cells with gaps was no longer significant from baseline levels by 1 week after noise (n.s., p=0.68). However, in *Xirp2* knockout mice, although we observed a smaller, non-significant (p=0.45) initial increase in cells with gaps after noise, the percent of cells with gaps actually further increased during the following week (**, p=0.003) (**Fig 4E**), rather than decreasing to pre-noise levels like in wild type, suggesting that gaps are not efficiently repaired in the absence of XIRP2. We also do not see a significant difference in the number of dead IHCs (**Fig. 4F**) or in stereocilia number (**Fig. 4G**) in wild type or *Xirp2* knockout mice at any timepoint following noise exposure, making it unlikely that hair cell death or stereocilia loss contribute to the observed differences.

### XIRP2 is necessary for the enrichment of monomeric γ-actin at gaps

We hypothesized that XIRP2 could be acting in a similar manner as zyxin and paxillin do at stress fibers (Smith et al., 2013, 2010), by recruiting factors necessary for the repair of stereocilia F-actin damage, including monomeric actin. As was previously shown (Belyantseva et al., 2009), *γ−*actin immunostaining is enriched in gaps in hair cell stereocilia (**Fig. 5A**). A representative plot of the fluorescence intensity of *γ*-actin staining shows a ∼2.7-fold increase at the center of the phalloidin-negative gap compared to the surrounding region (**Fig. 5B**). *γ−*actin at these sites is likely to be primarily monomeric, as the phalloidin signal is weak, and DNaseI staining, which labels monomeric *β*- and *γ*-actin (Mannherz et al., 1975), is also enriched (Belyantseva et al., 2009). However, in contrast to wild type mice, *γ*-actin no longer appears to be enriched in gaps in *Xirp2* knockouts (**Fig. 5C**). The plot profile in **Fig. 5D** shows the absence of *γ*-actin fluorescence intensity enrichment in an exemplary gap from a *Xirp2* knockout mouse. On average, the ratio of *γ*-actin fluorescence intensity at the center of phalloidin-negative gaps compared to the edge of gaps decreases from 2.03±0.16-fold in wild type mice to 1.29±0.06-fold in the absence of XIRP2 (***, p=0.0008) (**Fig. 5E**).

**Figure 5.**
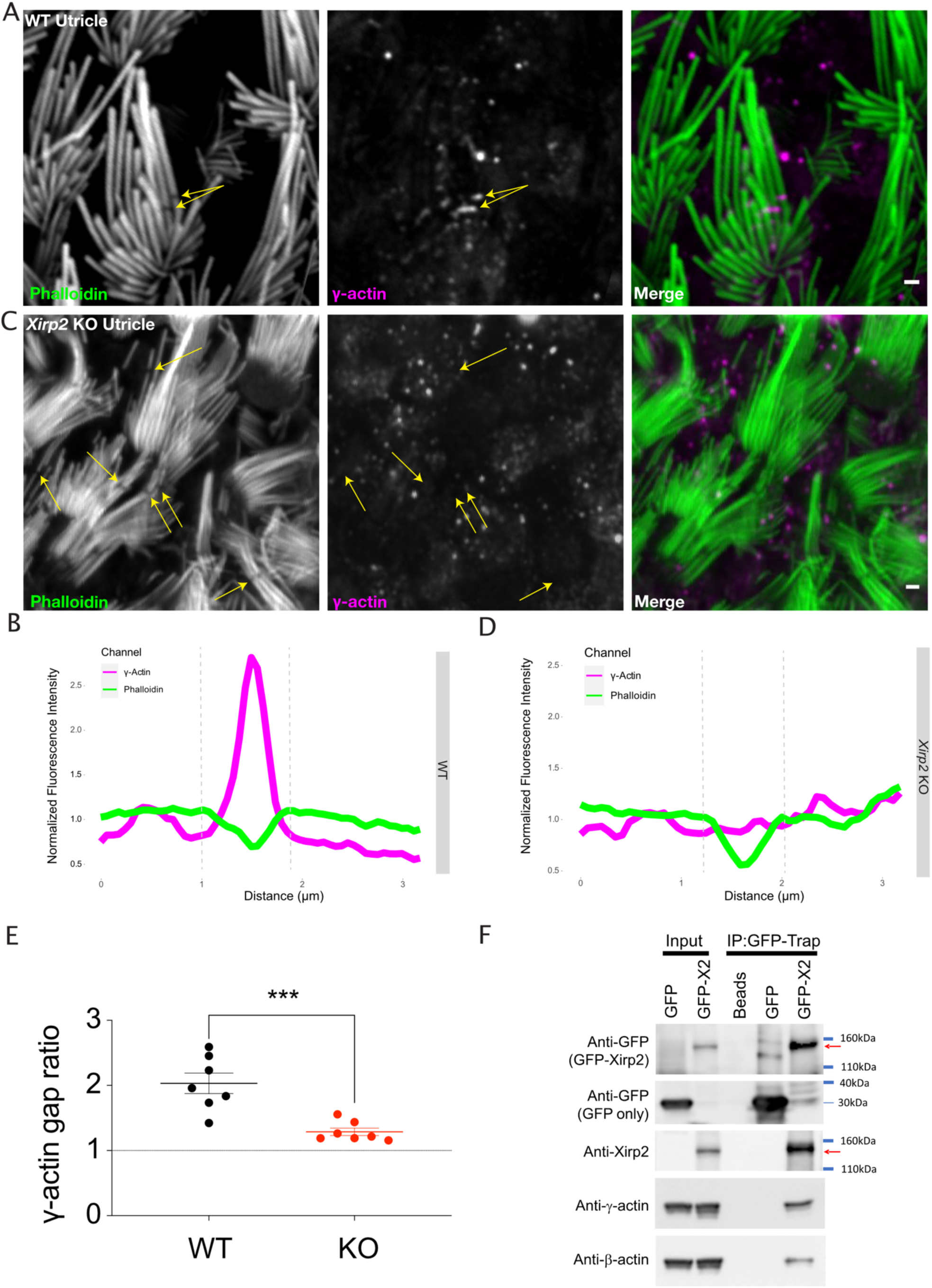
Monomeric γ-actin is no longer recruited to gaps in the absence of XIRP2. **(A)** *γ*- actin immunostaining is enriched in gaps (yellow arrows) in utricle hair cells. **(B)** Line scan of fluorescence intensity in phalloidin (green) and *γ*-actin (magenta) channels along the length of a gap in a WT mouse where *γ*-actin immunostaining is enriched. **(C)** *γ*-actin gap enrichment is decreased in *Xirp2* knockout mice (yellow arrows). **(D)** Line scan of fluorescence intensity in phalloidin (green) and *γ*-actin (magenta) channels along the length of a gap in a *Xirp2* knockout mouse where *γ*-actin immunostaining is enriched. **(E)** The enrichment of *γ*-actin at gaps is decreased in *Xirp2* knockout mice (***, p<0.001). **(F)** Monomeric *β*- and *γ*-actin co- immunoprecipitate with heterologously expressed short XIRP2. NIH 3T3 cells were transfected with the GFP or GFP-XIRP2 construct indicated on the top of each lane. Total cell extract was loaded in Input lanes. Immunoprecipitates (IP) were pulled down with GFP-Trap agarose beads followed by western blotting for the indicated antibodies (left). GFP-Trap beads (Beads) were incubated with non-transfected cell extracts as a negative control. All scale bars are 1μm. Error bars represent SEM. Images are representative of >3 experiments.

The lack of *γ*-actin enrichment at gaps in *Xirp2* knockout mice led us to hypothesize that short XIRP2 can interact directly or indirectly with actin. To address this, we co-immunoprecipitated endogenously-expressed *γ*- and *β*-actin in NIH-3T3 cells with GFP-tagged short XIRP2. To differentiate between monomeric and filamentous actin, we depolymerized the actin filaments in the cell lysate with MgCl_2_ (Wang et al., 2003). Depolymerized *γ*- and *β*-actin were both detected in the immunoprecipitate (**Fig. 5F**), suggesting that short XIRP2 interacts with both actin isoforms in their monomeric form, but does not exclude the possibility that it also interacts with F-actin.

### The C-terminus of the short isoform of XIRP2 is required for its role at gaps

Because some LIM domains have been shown to bind to tensed F-actin (Sun et al., 2020) and the LIM domains of zyxin and paxillin are necessary and sufficient for their recruitment to SFSS (Smith et al., 2013), we hypothesized that the C-terminal domain of the short isoform, which contains a predicted LIM domain, would be required for its recruitment to gaps. To test this possibility, using CRISPR/Cas9-mediated homology directed repair, we made a mouse in which the C-terminus of XIRP2 is removed. Due to the overlapping but differing reading frames in the C-termini long and short isoforms of XIRP2, it was impossible to specifically remove the LIM domain of short XIRP2 without affecting long XIRP2 (**Fig. 6A**). However, by making a 1 base pair substitution in exon 8 (T1187A, KM273012.1), we were able to introduce a stop codon before the beginning of the LIM domain in short XIRP2 (L396*, AIR76303.1) without altering the amino acid coding sequence of the long isoform (V3255V, NP_001077388.1) (**Fig. 6B**). This mutation led to a truncation of short XIRP2 in which the C-terminus (including the LIM domain) is removed (**Fig. 6C**). Truncated short XIRP2 (XIRP2-ΔCterm) is still expressed and localizes to the hair bundle, as XIRP2 staining with an antibody recognizing the N-terminus of both isoforms is still present in stereocilia (**Fig. 6D**). However, immunostaining with an antibody recognizing the truncated C-terminus is absent (**Fig. 6E**). To confirm that the bundle staining in the *Xirp2-*Δ*Cterm* mice was not due to compensatory relocalization of long XIRP2, we found that long XIRP2, recognized by an antibody against the alternative reading frame in exon 8, is still primarily localized to the cuticular plate and cell junctions (**Fig. 6F**).

**Figure 6.**
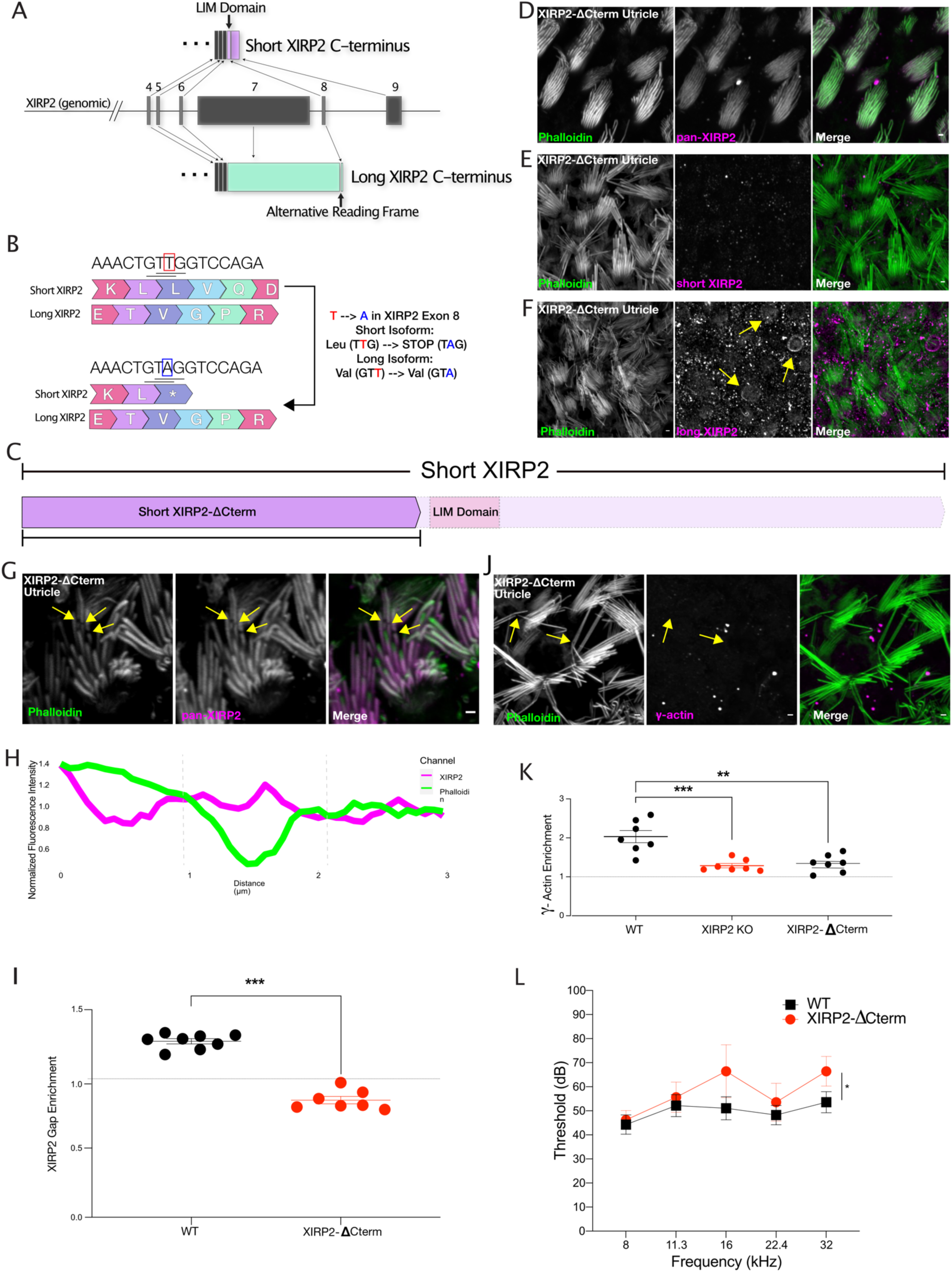
The C-terminus of XIRP2 is required for its recruitment to gaps. **(A)** Diagram of *Xirp2* gene structure and isoforms indicating the position of the LIM domain and the region of alternative reading frame targeted in the generation of the *Xirp2-*Δ*Cterm* mice. **(B)** Diagram indicating the position of the 1 bp substitution in exon 8 of *Xirp2* to generate the *Xirp2-*Δ*Cterm* mice. The TàA mutation introduced a stop codon in the short isoform but did not alter the amino acid sequence of the long isoform. **(C)** The LIM domain of short XIRP2 and the rest of the C- terminus is removed from short XIRP2 in *Xirp2-*Δ*Cterm* mice. **(D)** XIRP2-ΔCterm (recognized by an antibody against the N-terminus) is still expressed and localizes to the hair bundle. **(E)** Immunostaining with an antibody targeting the C-terminus of XIRP2 is absent in *Xirp2-*Δ*Cterm* mice, indicating the successful truncation of short XIRP2. **(F)** The bundle signal in *Xirp2-*Δ*Cterm* mice is not due to compensatory localization of long XIRP2. **(G)** Unlike full length short XIRP2, XIRP2-ΔCterm immunostaining is not enriched at gaps (yellow arrows). **(H)** Line scan of fluorescence intensity in phalloidin (green) and XIRP2 (magenta) channels along the length of a gap in *Xirp2-*Δ*Cterm* mice in which XIRP2 is not enriched. **(I)** The enrichment of XIRP2 immunostaining at gaps is decreased from 1.27-fold in WT mice to 0.85-fold in *Xirp2-*Δ*Cterm* mice (***, p<0.0.001). n= 8 utricles, 4 mice per group. **(J)** *γ*-actin immunostaining enrichment is decreased in gaps (yellow arrows) in *Xirp2-*Δ*Cterm* mice. **(K)** *γ*-actin gap enrichment is decreased from ∼2-fold in WT to 1.3-fold in *Xirp2-*Δ*Cterm* mice (**, p=0.002). n= 7 utricles, 4 mice per group. **(L)** *Xirp2-*Δ*Cterm* mice have elevated hearing thresholds compared to WT mice at 5 months of age, as measured by ABR (*, p=0.031). n= 14 WT mice, 8 *Xirp2-*Δ*Cterm* mice. All scale bars are 1μm. Error bars represent SEM. Images are representative of >3 experiments.

Despite XIRP2-ΔCterm’s colocalization with stereocilia actin cores, accordant with our hypothesis, XIRP2-ΔCterm is not enriched in phalloidin-negative gaps (**Fig. 6G**). Rather than the enrichment seen in wild type mice, the plot profile in **Fig. 6H** shows the depletion of XIRP2 immunostaining at the center of a phalloidin-negative gap. The enrichment of XIRP2 fluorescence intensity at the center of gaps decreases from 1.27±0.02-fold in wild type mice to 0.85±0.03-fold in *Xirp2-*Δ*Cterm* mice (***, p<0.0.0001) (**Fig. 6I**).

The lack of XIRP2 enrichment in gaps in *Xirp2-*Δ*Cterm* mice allowed us to ask whether the localization of XIRP2 to gaps was necessary for the subsequent enrichment of *γ*-actin. Although truncated short XIRP2 is present in undamaged regions of stereocilia in *Xirp2-*Δ*Cterm* mice, we observed a decrease in *γ*-actin enrichment in gaps (**Fig. 6J**). The average ratio of *γ*-actin enrichment is decreased to 1.34±0.09-fold in the *Xirp2-*Δ*Cterm* mice (**, p=0.002) (**Fig. 6K**), indicating that XIRP2 gap enrichment is necessary for subsequent recruitment of *γ*-actin.

The *Xirp2-*Δ*Cterm* mice also allowed us to ask whether the presence of XIRP2 in gaps is important for hearing function. Albeit not as severe as the phenotype in *Xirp2* knockout mice, *Xirp2-*Δ*Cterm* mice have increased hearing thresholds compared to wild type mice, as measured by auditory brainstem response (*, p=0.031) (**Fig. 6L**), suggesting that gap repair may be important for maintaining hearing sensitivity.

### *Xirp2* knockout mice are more vulnerable to age-related and noise-induced hearing loss

Our above data demonstrate that stereocilia F-actin gaps can be repaired in a XIRP2-dependent manner. We next asked whether the gap repair is important for the maintenance of hearing sensitivity in aging animals and animals exposed to noise.

We previously reported that *Xirp2* knockout mice, which were maintained on a C57Bl/6J background, develop high frequency hearing loss by 7-8 weeks of age (Francis et al., 2015). Here, we expanded those studies and show that by 11 weeks, the hearing loss worsens to affect all frequencies (**Fig. 7A**) (***, p<0.001), and progressively worsens overtime, as observed at 18 (**Fig. 7B**) (***, p<0.001) and 40 weeks (**Fig. 7C**) (***, p<0.001). Progressive hearing loss in mice maintained on the C57Bl/6J background is frequently exacerbated due to a mutation in the tip link protein *Cdh23*, which is harbored in this strain (Johnson et al., 1997). To separate the effect of this mutation on the worsening hearing in *Xirp2* knockout mice, we also measured hearing function in *Xirp2* knockout mice backcrossed for 10 generations to the CBA/J background. Although the hearing loss is less severe in CBA/J *Xirp2* knockout mice, and ABR thresholds are not significantly elevated at 11-12 weeks of age (**Fig. 7D**) (n.s., p=0.090), CBA/J *Xirp2* knockout mice develop hearing loss at all frequencies by 18 weeks (**Fig. 7E**) (***, p<0.001), which further progresses by 40 weeks (**Fig. 7F**).

**Figure 7.**
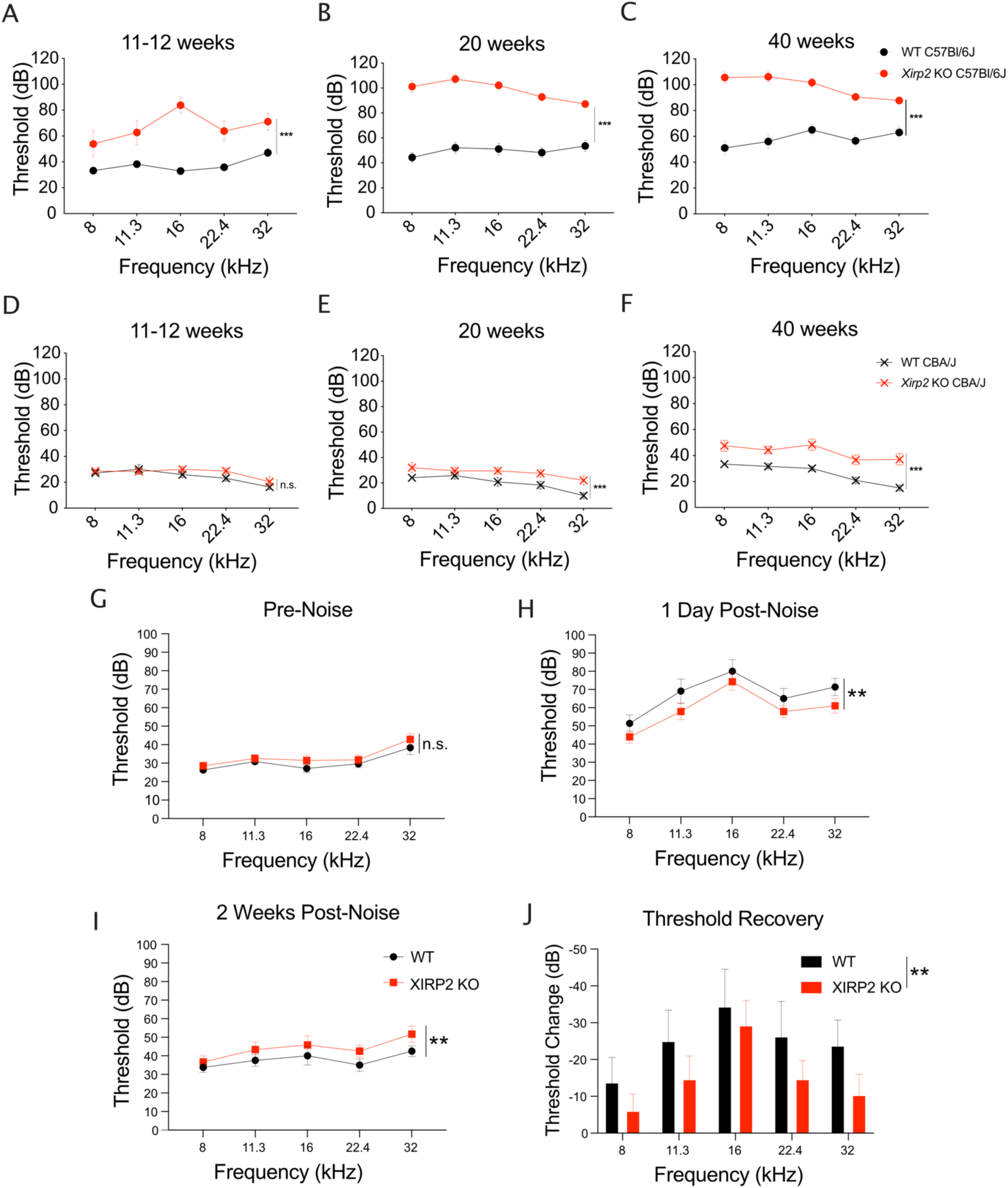
*Xirp2* knockout mice are more susceptible to age-related and noise-induced hearing loss. **(A)** ABR thresholds are significantly elevated in *Xirp2* KO mice on the C57Bl/6J background at 11-12 weeks of age (***, p<0.001). n=17 WT, 9 *Xirp2* KO. **(B)** ABR thresholds are further elevated in C57Bl/6J *XIrp2* KO mice at 18 weeks (***, p<0.001). n=14 WT, 9 *Xirp2* KO. **(C)** ABR thresholds are progressively increased in C57Bl/6J *Xirp2* KO mice at 40 weeks (***, p<0.001). n=10 WT, 9 *Xirp2* KO. **(D)** ABR thresholds are not significantly elevated in *Xirp2* KO mice on the CBA/J background at 11-12 weeks of age (n.s, p=0.090). n=11 WT, 12 *Xirp2* KO. **(E)** By 18 weeks, ABR thresholds are significantly elevated in CBA/J *Xirp2* KO mice (***, p=<0.001). n=6 WT, 12 *Xirp2* KO. **(F)** ABR thresholds are further increased in CBA/J *Xirp2* KO mice by 40 weeks of age (***, p=<0.001). n=6 WT, 12 *Xirp2* KO. **(G)** Prior to noise exposure, *Xirp2* knockout mice backcrossed to the CBA/J background do not have significantly different ABR hearing thresholds than WT CBA/J mice (n.s., p=0.058). n= 12 WT mice, 14 *Xirp2* KO mice. **(H)** Following a noise-induced temporary hearing threshold shift (1 day following noise exposure (105dB octave band, 1 hr)), WT CBA/J mice have elevated ABR hearing thresholds compared to XIRP2 KO CBA/J mice (**, p=0.007). n=12 WT mice, 14 *Xirp2* KO mice. **(I)** Following 2 weeks recovery, *Xirp2* KO CBA/J mice have elevated ABR hearing thresholds compared to WT CBA/J mice (**, p=0.009). n=8 WT mice, 14 *Xirp2* KO mice. **(J)** *Xirp2* KO CBA/J mice have decreased ABR hearing threshold recovery during the two weeks following exposure to noise. (**, p=0.005). n= 8 WT mice, 14 *Xirp2* KO mice. All error bars represent SEM.

We further hypothesized that repair of damaged stereocilia cores could contribute to the observed partial recovery of hearing thresholds following traumatic noise exposure (Wang et al., 2002). If this is the case, we would expect to see a similar initial temporary threshold shift (TTS) in wild type mice and the gap repair-deficient *Xirp2* knockout mice, but decreased recovery in *Xirp2* knockouts, leading to a larger permanent threshold shift (PTS). To test this possibility, we exposed wild type and *Xirp2* knockout mice on CBA/J background to PTS-inducing noise (105 dB for 30 minutes, 8-16kHz octave band (Paquette et al., 2016)) at an age (2 months) where there was no significant baseline hearing loss in CBA/J *Xirp2* knockout mice (**Fig. 7G**). Following the noise exposure, we surprisingly saw a smaller temporary threshold shift in the *Xirp2* knockout mice compared to wild types, as indicated by lower hearing thresholds 1 day following noise (**, p=0.007) (**Fig. 7H**). However, 2 weeks following the noise exposure, the *Xirp2* knockout mice display a larger PTS than wild types, with higher hearing thresholds across all measured frequencies at this time point (**, p=0.009) (**Fig. 7I**). For each individual mouse, we measured the change in threshold between 1 day and 2 weeks following noise exposure and found a decrease in recovery during this period in *Xirp2* knockout mice (**Fig. 7J**) ((**, p=0.005), accordant with our hypothesis that gap repair contributes to recovery from noise-induced hearing loss.

## Discussion

Acoustic trauma has been shown to cause disorganization and decreased negative staining of stereocilia actin filaments in TEM imaging (Engström et al., 1983; Liberman, 1987; Liberman and Dodds, 1987; Tilney et al., 1982) or gaps in phalloidin staining in confocal imaging (Avinash et al., 1993; Belyantseva et al., 2009). In agreement with this work, we found an increase in phalloidin-negative gaps following noise, and additionally observed gaps, albeit at a much lower rate, in auditory hair cells and vestibular hair cells in mice that were not exposed to damaging levels of noise, suggesting that the stereocilia cores may also be damaged from continuous exposure to environmental noise or head movement. These changes in actin visualization likely represent areas with decreased actin filament density, where actin filaments have been broken and depolymerized due to mechanical forces from overstimulation or changes in stereocilia ion concentrations from increased MET, as phalloidin binds at the interface of actin monomers within filaments (Barden et al., 1987; Cooper, 1987). Further studies will be needed to dissect the contributions of mechanical versus chemical contributions to the generation of noise-induced lesions in stereocilia cores.

Even in the absence of other hair bundle damage that occurs in concert with the stereocilia breakage during noise exposure (Wagner and Shin, 2019), we expect that gaps in the F-actin core decrease the structural integrity of the hair bundle and lead to decreases in MET function. Previous work has shown that *in vitro* fluid jet overstimulation of the hair bundle causes a decrease in bundle stiffness (Duncan and Saunders, 2000; Saunders et al., 1986; Saunders and Flock, 1986) without breakage of interstereociliary links (including the tip links), which was proposed to be due to damage to damage to the stereocilia core or to the rootlet where the stereocilia is inserted into the cuticular plate (Duncan and Saunders, 2000). Consistent with our observed recovery of stereocilia cores through actin remodeling, recovery of bundle stiffness was observed ((Duncan and Saunders, 2000; Saunders and Flock, 1986), dependent on hair cell metabolic activity (Saunders et al., 1986). The recovery occurred on a faster time scale than that which we observed, but it is possible that the intensity of damage was less severe due to the *in vitro* nature of the experiment or that recovery time is variable between species. Additionally, we cannot rule out repair of damage to the stereocilia rootlet, which was difficult to evaluate with confocal microscopy, contributed to stiffness recovery in their experiments. Further investigation of the actin core ultrastructure will be necessary to evaluate this possibility.

The vulnerability of stereocilia cores is compounded by the extremely slow turnover of F-actin. In many actin-based structures actin is continuously turned over by a process known as treadmilling, in which actin monomers are added to the plus-end of the filament at the same rate as they are removed from the minus end, maintaining the filament at a constant length (dos Remedios et al., 2003). In contrast, stereocilia actin is extremely stable, except for a small dynamic region at the very tips (Drummond et al., 2015; Narayanan et al., 2015; Zhang et al., 2012), preventing the damaged region from being replaced passively through turnover and necessitating an active repair mechanism to restore bundle rigidity and MET function. Actin monomers, an actin crosslinker, and an actin nucleator, which could serve as building blocks for new F-actin assembly, were found to be enriched at phalloidin negative gaps (Belyantseva et al., 2009), supporting this possibility. Combined with the decrease in gaps we found following noise exposure and incorporation of *β*-actin synthesized following noise exposure in discrete patches along the stereocilia shaft, we posit that these factors are recruited to gaps to facilitate the assembly of new actin polymers to replace the damaged filaments. Although we do observe newly synthesized *β*- actin in repaired gaps, it is possible that depolymerized actin from damaged filaments is also recycled during the repair process, which, together with the low Cre recombination rate, might explain the low frequency at which were able to observe repaired gaps harboring newly synthesized actin. As we did not observe GFP-labeled *β*-actin along the whole length of the stereocilia following noise, we propose that short filaments spanning the length of the gap are synthesized and attached to existing undamaged regions of the actin core through direct annealing, which has been described *in vitro* (Murphy et al., 1988). Although actin polymerization at the minus end of the filament is less energetically favorable than at the plus end, another possibility is that actin monomers are added on to filaments on both sides of the gap to fill in the damaged region and then the pieces are annealed together, possibly supported by the split staining of XIRP2 on either side of some gaps.

Actin monomers appear to be recruited to gaps in a XIRP2-dependent manner, as demonstrated by the decrease in *γ*-actin gap enrichment in *Xirp2* knockout mice. We also found that heterologously-expressed XIRP2 interacts with both monomeric *β*- and *γ*-actin and this interaction is presumably important for their recruitment to gaps. The decrease in monomeric actin gap localization is potentially the primary reason for the lack of gap repair that we observe in *Xirp2* knockout mice, as evidenced by increased gaps in *Xirp2* knockout mice in the absence of noise trauma and lack of decrease in gaps following noise exposure.

We previously reported that among the multiple XIRP2 isoforms expressed in hair cells, short XIRP2 (<u>Genbank ID</u> KM273012) specifically and exclusively localizes to the hair bundle (Francis et al., 2015). Immunostaining with isoform-specific antibodies demonstrated that only short XIRP2 is enriched at gaps in phalloidin staining. In contrast to the long XIRP2 isoform, short XIRP2 contains a C-terminal LIM domain. Our studies on the *Xirp2-*Δ*Cterm* mice where the C-terminus of short XIRP2, including the LIM domain is removed, suggest that this LIM domain mediates XIRP2 targeting to the F-actin lesion sites. This is consistent with reports of other LIM domain-containing proteins shown to bind specifically to tensed F-actin fibers, which may be present in gaps (Sun et al., 2020). Any remaining filaments within damaged regions are likely under increased stress from continued mechanical stimulation of the bundle. LIM domains have also been implicated in mediating interactions between many cytoskeleton-associated proteins (Kadrmas and Beckerle, 2004). The C-terminal LIM domain of XIRP2 might recognize another protein associated with the damaged stereocilia F-actin. Our study does not eliminate the possibility that the remaining portion of the truncated C-terminus is also important for gap recognition, but the lack of other predicted domains does not provide any mechanistic clues to suggest how it might be acting. It is also possible that the LIM domain has an additional role in recruiting actin nucleators and crosslinkers to gaps, similar to the LIM domains of zyxin and paxillin (Smith et al., 2013, 2010). Future work will address whether espin and cofilin are enriched at gaps in absence of the LIM domain of short XIRP2.

Interestingly, XIRP2 appears to remain at damaged sites after they have been repaired, as we see XIRP2 co-enriched with patches of newly synthesized *β*-actin-GFP at presumed sites of repair and the number of XIRP2 enriched sites following noise exposure remains elevated, despite the decrease in gaps during this time period. One possibility for XIRP2’s continued presence might be that gaps are not fully repaired over the 1 week period following noise exposure, but this is unlikely unless there is very minor damage remaining which cannot be observed with the resolution of light microscopy. However, if this is not the case, it may indicate that XIRP2 has a further role following the repair of the actin core, possibly in reenforcing the structure at vulnerable sites.

It is difficult to directly address the effects of gap formation and repair on hearing function because we do not have a method to specifically induce stereocilia core damage *in vivo* without damaging other hair cell structures. Much research on age-related and noise-induced hearing loss focuses on hair cell death and damage to the stria vascularis, which maintains the ion concentrations necessary for MET (Wu et al., 2020). However, there is evidence to suggest that unrepaired damage to the hair bundle, possibly including actin core lesions, also plays a role in the development of noise-induced hearing loss (Liberman and Dodds, 1984). The *Xirp2-*Δ*Cterm* mice allowed us to partially separate XIRP2’s function in gaps from its other, as of yet uncharacterized, roles in hair cells. Truncated short XIRP2 is still expressed in these mice and still appears to be localized normally in the hair bundle, except it is no longer enriched in gaps. If, as we suspect, XIRP2-ΔCterm is still able to perform the other functions of XIRP2, the hearing loss in *Xirp2-*Δ*Cterm* mice suggests that lack of gap repair leads to the development of hearing loss in those mice, even in the absence of noise trauma. The connection between repair of gaps and hearing loss is also supported by the progressive increase in hearing thresholds in *Xirp2* knockout mice. The buildup of unrepaired gaps in the absence of XIRP2 may lead to the worsening phenotype, suggesting that gap repair could be necessary for the prevention of age-related hearing loss.

We also propose here that repair of gaps may contribute to partial recovery from noise-induced hearing loss. Repair of other hair bundle structures like tip links (Indzhykulian et al., 2013; Zhao et al., 1996), which are broken following noise exposure (Husbands et al., 1999; Pickles et al., 1987), has been hypothesized to contribute to the decrease in hearing thresholds following an initial large temporary threshold shift. The increased susceptibility of *Xirp2* knockout mice to permanent noise-induced hearing loss is consistent with actin core repair also contributing to this threshold recovery. Although the initial temporary threshold shift was unexpectedly smaller in *Xirp2* knockout mice compared to wild type, the recovery over the 2-week period following noise exposure was dampened in *Xirp2* knockouts, leading to mildly, but significantly, elevated hearing thresholds. The relatively small difference between genotypes was unsurprising, however, if repair of gaps is only one of many factors contributing to this recovery, most of which we do not predict to be altered in the absence of XIRP2.

This study sheds further light on the importance of repair of sublethal damage to hair cells in order to preserve hearing function following traumatic noise exposure and in aging animals. Due to the inability of mature mammalian hair cells to regenerate (Burns and Corwin, 2013; Groves, 2010), repair mechanisms are necessary to restore mechanotransduction and prevent hearing loss. Herein, we add stereocilia actin cores to the growing list of hair cell structures, including tip links (Indzhykulian et al., 2013) and, controversially, synapses (Kujawa and Liberman, 2009; Shi et al., 2013), capable of recovery in mammalian hair cells.

## Methods

### Animal care and handling

The care and use of animals for all experiments described conformed to NIH guidelines. Experimental mice were housed with a 12:12 h light:dark cycle with free access to chow and water, in standard laboratory cages located in a temperature and humidity-controlled vivarium. The protocols for care and use of animals was approved by the Institutional Animal Care and Use Committees at the University of Virginia, which is accredited by the American Association for the Accreditation of Laboratory Animal Care. C57BL/6J (Bl6, from Jackson Laboratory, ME, USA) and CBA/J (from Jackson Laboratory, ME, USA) mice and sibling mice served as control mice for this study. Neonatal mouse pups (postnatal day 0 (P0)–P5) were sacrificed by rapid decapitation, and mature mice were euthanized by CO_2_ asphyxiation followed by cervical dislocation. The *FLEx-β-actin-GFP* mice were purchased from The Jackson Laboratory. The *Ubc-Cre^ERT2^* mice were gifted from Dr. Alban Gaultier (University of Virginia).

All procedures involving *Elmod1 knockout* animals met the guidelines described in the National Institutes of Health Guide for the Care and Use of Laboratory Animals and was approved by the Animal Care and Use Committees of Tel Aviv University (M-12-046).

### Generation of *Ptprq* knockout and *Xirp2-*Δ*Cterm* mice

For CRISPR/Cas9 mediated generation of the mouse models, we used the CRISPR design bioinformatics tool developed by Feng Zhang’s lab at the Massachusetts Institute of Technology (crispr.mit.edu (Hsu et al., 2014)) and the online tool CRISPOR (http://crispor.tefor.net/crispor.py) to select suitable target sequences to knockout *Ptprq* (CTCACCCGTGAGTAGAACAC, in exon 2) and to edit the sequence in XIRP2 to generate the *Xirp2-*Δ*Cterm* mice (GAAACTGTTGGTCCAAGACA, in exon 8 and the preceding intron), respectively. The following repair template was used to generate the 1 bp substitution in exon 8 to introduce a stop codon in the short isoform: CCAGTTAGTGTAGTATTTGTTTTGTTATGCCTTTACAGTTGTGACTGAAAATAGAAATATATA AGCCTTTATATATATTTTTTTATTTTTAGAAACTGT**<u>A</u>**GGTCCAAGGCAAGGAAATTTGCATAA TTTGTCAAAAGACAGTTTATCCAATGGAGTGCCTCATAGCAGACAAGCAGAATTTTCATAAGTCTTGCTTCAGA (1bp TàA substitution indicated by bolded A). The repair template was ordered as a custom Ultramer DNA oligo from Integrated DNA Technologies. The single-guide (sg)RNA to target exon 8 of *Xirp2* was generated by T7 *in vitro* transcription of a PCR product from overlapping oligonucleotides (as described in the CRISPOR online tool). Oligonucleotide sequence for the *Xirp2-*Δ*Cterm* knock-in forward primer: GAAATTAATACGACTCACTATAGGAAACTGTTGGTCCAAGACAGTTTTAGAGCTAGAAATAG CAAG, and for the universal reverse primer: AAAAGCACCGACTCGGTGCCACTTTTTCAAGTTGATAACGGACTAGCCTTATTTTAACTTGC TATTTCTAGCTCTAAAAC. The sgRNA to target exon 2 of *Ptrpq* was generated by *in vitro* transcription of the target sequence cloned downstream of the T7 promoter in the pX330 vector (Addgene). *In vitro* transcription was performed using the MAXIscript T7 kit (Life Technologies) and RNA was purified using the MEGAclear kit (Thermo Fisher Scientific). For production of genetically engineered mice, fertilized eggs were injected with precomplexed RNP (ribonucleoprotein)-Cas9 protein (PNA Bio, 50 ng/μl) and sgRNA (30 ng/μl). In the case of the *Xirp2-ΔCterm* mice, the repair template was co-injected (50 ng/μl). Two-cell stage embryos were implanted on the following day into the oviducts of pseudopregnant ICR female mice (Envigo). Genotyping was performed by PCR amplification of the region of interest (*Ptprq* forward primer: ACTTTGGCATTCCAGGTTGATGT, *Ptprq* reverse primer: ATGCAAAGCAAACTCGGCCAAT; *Xirp2-*Δ*Cterm* forward primer: AGGCTCCCCATAAGTTTGCT, *Xirp2*-ΔCterm reverse primer: TGCTTGTCTGCTATGAGGCA), followed by Sanger sequencing. Founder mice were selected and backcrossed for 7 generations to the C57Bl/6J background. The PTPRQ knockout mouse harbors a reading frame-shifting 1-bp deletion in exon 2. The *Xirp2-*Δ*Cterm* mice have the intended 1bp substitution with no other edits.

### Generation of *Elmod1* knockout mice

The ES cells of *Elmod1*-deficient mice (Elmod1^tm1a (EUCOMM)Hmgu^) were produced at the EUCOMM Consortium and the mice were produced at MRC Harwell. Mice carrying a cassette including LacZ and neomycin resistance genes inserted into intron 4-5 of the *Elmod1* gene, located on chromosome 9. The EMMA ID is EM:04834, common strain name: HEPD0557_4_E10, and international strain designation: C57BL/6N-Elmod1^tm1a(EUCOMM)Hmgu^/H, are referred to herein as *Elmod1* knockout. The mice were maintained on a C57BL/6NTac genetic background and were bred with wild-type C57BL/6. For genotyping, genomic DNA was extracted from a small clip of the tail and was used as a template for PCR with specific primers. For the wild-type allele the primers used were: forward, GGGGTGACCTAAGACCATCA; and reverse, GAGGGCTCGGGAGACATAGT. For the mutant allele: forward, GTTTGTTCCCACGGAGAATC; reverse, CCGCCACATATCCTGATCTT. The mutant line is available from the Infrafrontier Mouse Disease Models Research Infrastructure (https://www.infrafrontier.eu/).

### Hearing tests in mice

ABRs of C57Bl/6J WT, CBA/J WT, *Xirp2* knockout C57Bl/6J, *Xirp2* knockout CBA/J, and *Xirp2-*Δ*Cterm* mice were determined at the time point indicated for longitudinal ABRs. Additionally, ABRs of CBA/J WT and *Xirp2* knockout mice were determined at 5-6 weeks on the day prior to and after noise exposure. An additional ABR was performed on the same mice at 2 weeks following noise exposure. Mice were anesthetized with a single intraperitoneal injection of 100 mg/kg ketamine hydrochloride (Fort Dodge Animal Health) and 10 mg/kg xylazine hydrochloride (Lloyd Laboratories). ABRs were performed in a sound-attenuating cubicle (Med-Associates, product number: ENV-022MD-WF), and mice were kept on a Deltaphase isothermal heating pad (Braintree Scientific) to maintain body temperature. ABR recording equipment was purchased from Intelligent Hearing Systems (Miami, Fl). Recordings were captured by subdermal needle electrodes (FE-7; Grass Technologies). The noninverting electrode was placed at the vertex of the midline, the inverting electrode over the mastoid of the right ear, and the ground electrode on the upper thigh. Stimulus tones (pure tones) were presented at a rate of 21.1/s through a high-frequency transducer (Intelligent Hearing Systems). Responses were filtered at 300–3000 Hz and threshold levels were determined from 1024 stimulus presentations at 8, 11.3, 16, 22.4, and 32 kHz. Stimulus intensity was decreased in 5–10 dB steps until a response waveform could no longer be identified. Stimulus intensity was then increased in 5 dB steps until a waveform could again be identified. If a waveform could not be identified at the maximum output of the transducer, a value of 5 dB was added to the maximum output as the threshold.

### Human utricle preparation

Human utricles used in this study were obtained as a byproduct of a surgical labyrinthectomy. Patient information was de-identified. The samples were fixed immediately after removal in a 10% formalin solution.

### Immunofluorescence

Inner ear organs were fixed in 4% paraformaldehyde (PFA, Electron Microscopy Sciences) immediately after dissection for 20 min. Samples were washed three times with phosphate-buffered saline (GIBCO^®^ PBS, Thermo Fisher Scientific) for 5 min each. After blocking for 2 h with blocking buffer (1% bovine serum albumin (BSA), 3% normal donkey serum, and 0.2% saponin in PBS), tissues were incubated in blocking buffer containing primary antibody at 4 °C overnight. Samples were washed again three times for 5 minutes with PBS and then incubated in blocking buffer containing secondary antibody at RT for 1.5 hrs. Following secondary antibody incubation samples were washed 3 times for 5 minutes with PBS and then mounted on glass slides in ProLong Glass Antifade Mountant (P36984, Invitrogen) with a #1 coverslip. The following primary antibodies were used in this study: goat polyclonal pan-XIRP2 antibody used in Fig. 2A-B (D18 (discontinued), Santa Cruz Biotechnology, Inc., 1:200), rabbit polyclonal pan-XIRP2 antibody used in Fig. 2C-D, Fig. 3, and Fig. 6 (custom generated against the immunizing peptide MARYQAAVSRGDTRSFSANVMEESDVCTVPGGLAKMKRQFEKDKMTSTCNAFSEYQYRHESRAEQEAIHSSQEIIRRNEQEVSKGHGTDVFKAEMMSHLEKHTEETKQASQFHQYVQETVIDSPEEEELPKVSTKILKEQFEKSAQENFLRSDKETSPPAKCMKKLLVQDKEICIICQKTVYPMECLIADKQNFHKSCFRCHHCSSKL, ProteinTech, 1:100), mouse monoclonal IgG2a GFP antibody (A11120, Invitrogen, 1:300), rabbit polyclonal long XIRP2 antibody (11896-1-AP, ProteinTech, 1:100), rabbit polyclonal short XIRP2 antibody (custom generated against the immunizing peptide NSKRQDNDLRKWGD, Genscript, 1:100), mouse monoclonal IgG_1_ *γ*-actin antibody (1-24: sc- 65635, Santa Cruz Biotechnology, Inc., 1:100). Alexa Fluor-488, -555, and -647 conjugated donkey anti-mouse IgG, -goat IgG, and -rabbit IgG secondary antibodies (Invitrogen, 1:200) were used to detect appropriately paired primary antibodies. F-actin was detected using Alexa Fluor- 488 and -647-conjugated Phalloidin (Invitrogen, 1:200). Fluorescence imaging was performed using an inverted Zeiss LSM880 confocal microscope with a fast-mode Airyscan detector and a 63x /1.4 NA, oil-immersion objective or an inverted Olympus FV1200 confocal microscope equipped with a 60x/1.35 NA oil-immersion objective.

### Image denoising

Denoising of confocal images was performed using the Noise2Void package, which is based on deep learning. The detailed mathematical model and network architecture for Noise2Void was previously published by the Jug group (Krull et al., 2019) and the package can be downloaded at https://github.com/juglab/n2v. An Intel i9-10900KF CPU and a single NVIDIA RTX3080 GPU were used for transfer learning. For each of the raw images, we trained a unique Noise2Void convolutional neural network model. In the transfer learning process, raw images of 1000×1000 pixels in x-y dimension were split into multiple 48×48 pixel tiles to make a training dataset. For training of each model, the network was trained with 200 epochs and 100 steps in each epoch. Finally, using the raw image as input, a denoised image can be predicted by using the trained neural network.

### Induction of Cre recombination

To induce the translocation of Cre to the nucleus in *FLEx β-actin-GFP+;Ubc-Cre^ERT2^+* mice, mice were intraperitoneally injected with tamoxifen. Two injections were performed 24 hours apart. Tamoxifen (T5648, Sigma) was dissolved at 5mg/ml in corn oil (C8267, Sigma) at 37°C for 2 hrs. Mice were injected with 9mg tamoxifen per 40 grams body weight.

### Noise exposure

Mice were placed separately in wire cages inside a custom-built small wooden reverberant box, built with instructions from the Charles Lieberman lab (Liberman and Gao, 1995), equipped with a high-intensity speaker (Model #2446H JBL Incorporated) and amplifier (Crown). To induce maximal stereocilia core damage for experiments in which gaps were quantified or newly synthesized actin incorporation localization was analyzed, mice were exposed to 4 hrs of broadband (1-22kHz) noise at 120 dB. Following noise exposure mice were allowed to recover for 1 hr, 8hrs, or 1 week. To induce a mild permanent hearing threshold shift (Paquette et al., 2016), mice were exposed to 105 dB octave band (8-16kHz) noise for 1 hr. ABRs were assessed 1 day before,1 day after, and 2 weeks after noise exposure.

### Gap quantification

For each experiment in which gaps were quantified in auditory hair cells, 3 images were collected for each cochlea at specified positions in the cochlear apex (5%, 20%, and 40% of full cochlear length). Each image contained ∼15 IHCs and the number of gaps in tallest row IHC stereocilia and cell number were quantified. The percent of cells with gaps was calculated for each cochlea. For experiments in which gaps were quantified in vestibular hair cells, 4 images were collected for each utricle (2 in the striola, 1 in the medial extrastriola, and 1 in the lateral extrastriola). Each extrastriolar image contained ∼80 HCs and each striolar image contained ∼60 HCs. The number of gaps and the cell number was quantified for each image and the percent of cells with gaps was quantified for each utricle.

### Cell death and stereocilia number quantification

Using the same images collected for quantification of gaps following noise exposure (3 images per cochlea at defined apical regions), the number of dead IHCs were quantified to calculate the percentage of dead IHCs per cochlea. The same images were also used to quantify changes in stereocilia number following noise exposure. The number of tallest row stereocilia for each IHC was quantified, ensuring that the counted stereocilia were attached to the cuticular plate at the rootlet. The average number of stereocilia per IHC was calculated for each cochlea.

### Quantification of gap enrichment immunostaining

Both utricles were collected from each of 4 mice and 4 images were taken from each utricle (2 striolar, 1 medial extrastriolar, and 1 lateral extrastriolar). For each phalloidin negative gap, a line scan of fluorescence intensity was generated for the phalloidin and XIRP2 or *γ*-actin channel. At the center of the gap (defined by the point with lowest phalloidin fluorescence intensity) and the edge of the gap (defined as the point where phalloidin signal begins to decrease) the fluorescence intensity was determined for XIRP2 or *γ*-actin. The intensity at the center of the gap was divided by the intensity at the edge of the gap to get the enrichment ratio. The average XIRP2 or *γ*-actin gap enrichment ratio was calculated for each utricle.

### Statistical analysis

For statistical analysis, GraphPad Prism (La Jolla, CA) and Microsoft Excel were used. Two-way ANOVA was used to determine statistically significant differences in the ABR and threshold changes following noise exposure recovery analyses. Two-tailed, unpaired Student’s *t* tests were used for other comparisons between two groups. *p* values smaller than 0.05 were considered statistically significant, other values were considered not significant (n.s.). Asterisks in the figures indicate *p* values (\**p* < 0.05, \*\**p* < 0.01, and \*\*\**p* < 0.001). All error bars indicate standard error of the mean (SEM). Sample sizes were determined based on variance from previous experiments in the lab for ABRs. For other experiments, sample size was determined from variance observed from pilot experiments. Whenever possible, quantifications were performed in a blinded manner.

### Co-Immunoprecipitation

NIH-3T3 cells were seeded on 6-well plates and allowed to grow to 70% confluency at 37°C and 5% CO_2_ in DMEM (Gibco) supplemented with 10% FBS (Life Technologies), 1 mm sodium pyruvate (Life Technologies), 100 U/ml penicillin, and 100 μg/ml streptomycin (Life Technologies). Media was exchanged with Opti-MEM (Gibco) and cells were transfected with either pCMV-GFP or pEGFP-XIRP2 using Lipofectamine 3000 (Invitrogen) according to manufacturer protocols. Following 48 hours incubation, cells were lysed with NP-40 lysis buffer (50mM Tris-HCl pH 8, 150mM NaCl, 1% NP-40, 5% glycerol, 2mM MgCl_2_, 2mM NaF, 1mM Na_3_VO_4,_ 1mM PMSF (Thermo Scientific), and Pierce protease inhibitor cocktail (Thermo Scientific)) on ice for 10 minutes. Cell lysates were sonicated (550 Sonic Dismembrator, Fisher Scientific) for 2 min (with one 10 sec pulse at amplitude 5 per 15 sec) and centrifuged at 13,000x*g* for 5 min. Cell lysates were added to GFP-Trap agarose beads (ChromoTek) pre-equilibrated with lysis buffer and incubated for 2hrs at 4°C. Beads were washed 3 times with lysis buffer with 30mM MgCl_2_ to depolymerize actin (Wang et al., 2003). 30 ul of 2X NuPAGE LDS sample buffer (Novex by Life Technologies) was added to the samples. 10 ul of each sample and 3% input of each cell lysate was loaded on a 4-12% Bis-Tris pre-cast SDS-page gel (Invitrogen) and transferred to a PVDF membrane (BioRad). The membrane was blocked with Odyssey Blocking Buffer (LI-COR) for 30 min at RT and incubated with the following primary antibodies at 4°C overnight in TBST with 5% BSA: monoclonal mouse GFP antibody (B-2: sc-9996, Santa Cruz Biotechnology, Inc., 1:500), polyclonal pan-XIRP2 antibody (custom generated (see above), ProteinTech, 1:1000), monoclonal mouse IgG_1_ *γ*-actin antibody (1-24: sc-65635, Santa Cruz Biotechnology, Inc., 1:1000), and mouse monoclonal *β*-actin antibody (AC-15: sc-69879, Santa Cruz Biotechnology, Inc., 1:1000). Membranes were washed 3 times for 5 min with TBST and then incubated with the corresponding secondary antibodies (donkey anti-mouse IgG or donkey anti-rabbit IgG conjugated HRP (Jackson ImmunoResearch) for 1hr RT at 1:10000 in TBST with 5% non-fat milk. Membranes were then washed 3 times for 4 min with TBST and treated with Pierce ECL Western Blotting Substrate (Thermo Scientific) according to manufacturer protocol. Membranes were imaged using the ImageQuant LAS system (GE Science).

## Acknowledgements

We would like to thank Alban Gaultier for providing the *Ubc-Cre^ERT2^* mouse line and Marissa Gonzales and Dr. Timothy Bullock for providing the NIH-3T3 cells. This study was supported by NIH/NIDCD grants R01DC014254, R56DC017724, and R01DC018842 (to J.-B.S.), R01DC011835 (to K.B.A.), and 1F31DC017370-01 (to E.L.W).

